# Supra-molecular organization of the pyoverdine bio-synthetic pathway in *Pseudomonas aeruginosa*

**DOI:** 10.1101/438713

**Authors:** Véronique Gasser, Morgane Malrieu, Anne Forster, Yves Mély, Isabelle J Schalk, Julien Godet

## Abstract

The bio-synthesis of the pyoverdine siderophore (PVD) in *Pseudomonas aeruginosa* involves multiple enzymatic steps including the action of Non-Ribosomal Peptide Synthetases (NRPS). One hallmark of NRPS is their ability to make usage of non-proteinogenic amino-acids synthesised by co-expressed accessory enzymes. It is generally accepted that different enzymes of a secondary metabolic pathway must organise into macro-molecular complexes. However, evidence for complexes like siderosomes in the cellular context are missing.

Here, we used *in vitro* single-molecule tracking and FRET–FLIM (Förster resonance energy transfer measured by fluorescence lifetime microscopy) to explore the spatial partitioning of the ornithine hydroxylase PvdA and its interactions with NRPS. We found PvdA was mostly diffusing bound to large complexes in the cytoplasm with a small exchangeable trapped fraction. FRET-FLIM clearly showed PvdA is physically interacting with PvdJ, PvdI, PvdL and PvdD, the four NRPS of the pyoverdine pathway. The binding modes of PvdA are strikingly different according to the NRPS it is interacting with suggesting that PvdA binding sites have co-evolved with the enzymatic active sites of NRPS.

Our data provide evidence for strongly organised multi-enzymatic complexes responsible for the bio-synthesis of PVD and suggest that finely controlled co-localisation of sequential enzymes seems to be required to promote metabolic efficiency.

## Introduction

To improve fitness, bacteria and fungi have evolved sophisticated machinery to produce secondary metabolites with remarkable structural and functional diversities. Secondary metabolites are often produced by specific and dedicated biosynthetic pathways activated in response to environmental stimuli. If much is understood about the enzymatic cascades and mechanisms that underlie these bio-syntheses, comparatively little is known about the cellular organisation of their components(1). There is a long-standing hypothesis that enzymes involved in a metabolic pathway must organise into macro-molecular complexes (2), so that secondary metabolite synthesis is orchestrated by the assembly of sequential enzymes. This process is expected to be widespread (3) (including in eukaryotic cells (4, 5)) and to play an important role in the overall bio-synthesis efficiency or regulation (5, 6). However observations on such specific multi-protein organisations are sparse in the cellular context and most of them are resulting from microscopy co-localisation observations (2).

Here we used the pyoverdine (PVD) metabolic pathway of *Pseudomonas aeruginosa* as a model system. PVDs are large fluorescent siderophores composed by an invariant (1S)-5-amino-2,3-dihydro-8,9-dihydroxy-1H-pyrimido[1,2-a]quinoline core grafted to a 6-to 12-amino acids chain (7, 8). In PAO1, PVD synthesis relies on four non-ribosomal peptide synthetases (NRPS)(9), PvdL, PvdI, PvdJ and PvdD (9–13) (Figure 1). NRPS are large modular enzymes synthesizing peptides independently from ribosomes (14, 15). Each NRPS module is responsible for the incorporation of a defined monomer into the final product (14) (Figure 1 C).The PVD precursor in PAO1 is an 11 amino-acid (AA) peptide with the sequence L-Glu – L-Tyr – D-Dab – L-Ser – L-Arg – L-Ser – L-fOHOrn – L-Lys – L-fOHOrn – L-Thr – L-Thr. The second and third amino acids of the peptide (L-Tyr and D-amino butyric acid) will form the chromophore in the PVD mature form. The use of non-proteinogenic amino acids, including D-isomers, is the hallmark of NRPS and is responsible for the sequence diversity of non-ribosomal peptides and derived products (16). Accessory and tailoring enzymes are required to synthesize modified amino-acids assembled by NRPS into the final peptide. In PAO1, PvdA (17), PvdF(18) and PvdH(13, 19) are involved in the synthesis of L-fOHOrn and D-Dab. Their genes are found at proximity of NRPS genes in PAO1 genome (Figure 1 B) in large bio-synthetic gene clusters (BGCs) (20, 21). These enzymes are thus likely co-expressed with NRPS when the PVD pathway is activated. Tailoring enzymes are also thought to impact critical protein–protein interactions on the main assembly line (22).

**Fig. 1.**
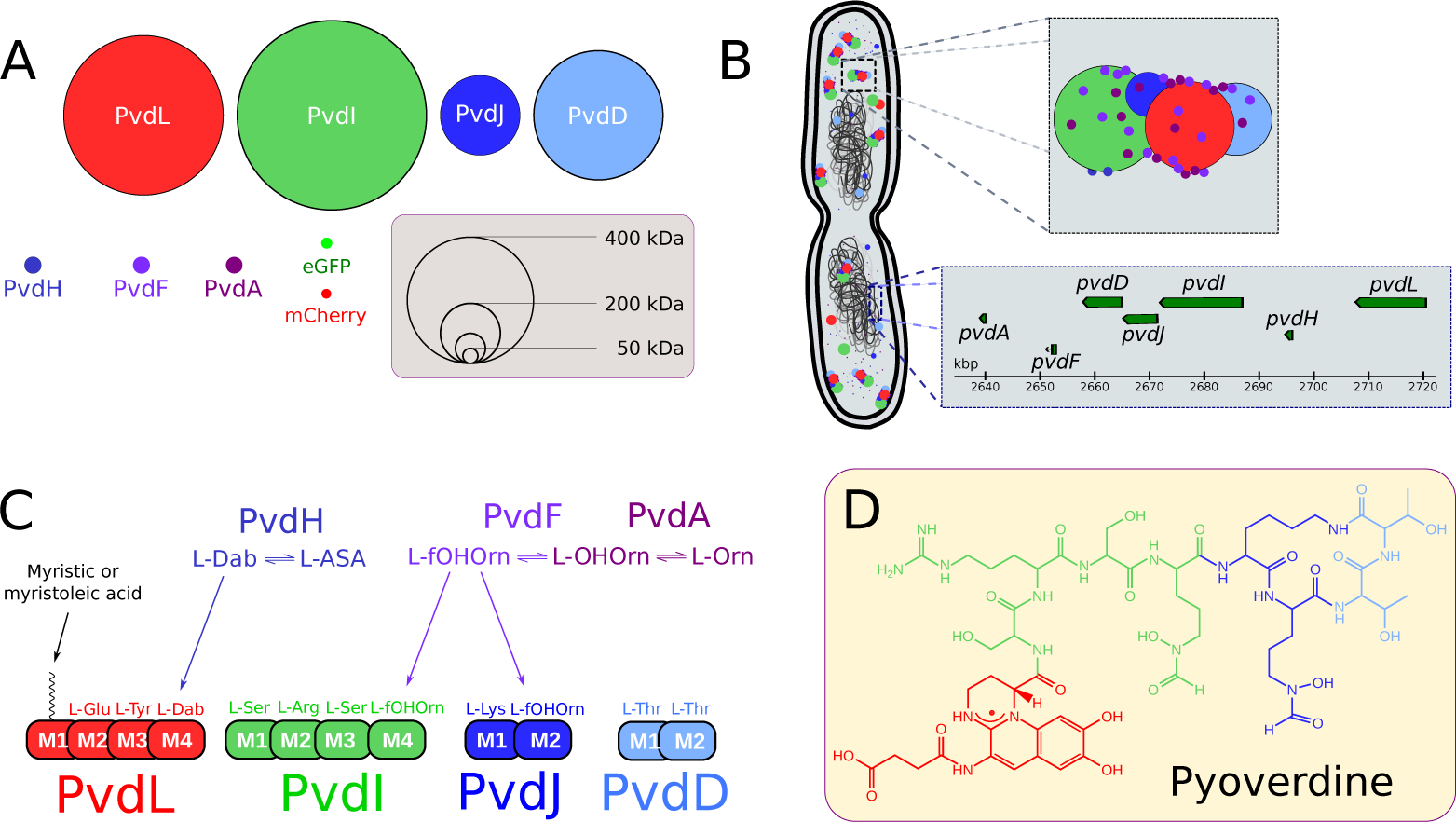
NRPS of *P. aeruginosa*. (A) Enzymes involved in the cytoplasmic bio-synthesis of the pyoverdine precursor in PAO1. Four large NRPS (PvdL, PvdI, PvdJ and PvdD) are responsible for the synthesis of a 11 amino-acid (AA) peptide and for the incorporation of modified AA, produced by PvdH(L-Dab) and by PvdA and PvdF (L-fOH Orn) accessory proteins. Enzyme diameters are proportional to their molecular weights (MW). eGFP and mCherry are shown for MW comparison. (B) Schematic view of the chromosomic genes and cytoplasmic proteins involved in the early steps of the pyoverdine bio-synthetic pathway in PAO1. Genes coding for NRPSs and accessory proteins are found in a large bio-synthetic gene cluster (BGC). Once the pyoverdine pathway is activated, proteins encoded by genes in the BGC are expressed and may organise in siderophore-specific multi-enzymatic complexes called siderosomes. (C) Pyoverdine precursor sequence. During bio-synthesis, each NRPSs module is responsible for the incorporation of a defined AA. PvdL (red), PvdI (green), PvdJ (blue) and PvdD (light blue) are acting sequentially to add respectively 3,4,2 and 2 AA of the 11-AA sequence of the pyoverdine precursor. (D) Pyoverdine (type I) - the major siderophore synthesised by PAO1 - is a maturated form of the cytoplasmic precursor.

PvdA catalyses the FAD-dependent hydroxylation of the amine side chain of ornithine using NADPH as the electron donor and molecular oxygen to synthesize N5-hydroxy-ornithine (23, 24). Recent fluorescence microscopy findings have shown a high cellular organization of PVD-related proteins with clusters of PvdA co-localising with PvdD, PvdL and PvdJ in fluorescence microscopy (25, 26). Pull-down assays using a recombinant 6His-PvdA protein as a bait captured low amount of PvdJ and PvdL (26). PvdA has also been shown to interact with PvdJ M2 isolated module in yeast double-hybrid experiment (26). Taken together, these data suggest that PvdA is interacting with NRPS and may be part of large multi-enzymatic complexes named siderosomes (25–27). However a clear biochemical characterisation of these complexes is missing – possibly due to highly dynamic interactions between the proteins of the siderosomes and/or their transient formation (27). Moreover only few attempts for characterising these complexes in the cellular environment have been reported. In this context, *in cellulo* characterisation with high temporal and/or spatial resolution are required to capture siderosomes organisation.

We investigated the localisation and dynamics of PvdA in cells using single-particle tracking combined with single-molecule localisation using photo-activated localisation microscopy (sptPALM (28)). The statistical description of thousands of single PvdA traces in cells showed that PvdA was mostly diffusing as large complexes in the cytoplasm, except for a small restricted and exchangeable fraction diffusing with a lower diffusion rate. We additionally used FRET measured by fluorescence lifetime imaging (FLIM (29)) to evidence PvdA interactions with PvdJ, PvdI, PvdL and PvdD. Although the exact determination of the complex stoichiometry was not accessible from FLIM-FRET measurements, our data are consistent with transient interactions of several PvdA on PvdI, PvdD and PvdL whereas PvdJ interacts with PvdA in a 1:1 stoichiometry. Our data evidence a complex organisation of siderosomes and suggest that PvdA binding sites have co-evolved with NRPS module specificity.

## Results

### PvdA diffuses within large complexes in the cytoplasm

To localise PvdA enzyme in its intracellular environment, we produced strains expressing at the chromosomal level PvdA fused to eGFP, eYFP or PAmCherry at its C-terminus as previously described (25, 27). These strains genetically expressed fluorescent PvdA at physiological levels. The expression of fluorescent PvdA was increased in iron-deprived growing conditions. These strains were able to produce and uptake PVD (see supplementary Figure S1). Epifluorescence microscopy showed that PvdA-eGFP was localised in the cytoplasm. The level of expression of PvdA-eGFP was heterogeneous from cell to cell. No spatial organisation was observed along the cell cycle, except for some unilateral polar spots in few cells (Figure 2 A-B). However, the resolution of epifluorescence microscopy is too low to rule out sub-diffraction limit (250 nm) organisations. To explore more in details the localisation and diffusion of single PvdA in live cells, we used Photoactivated Localisation Microscopy (PALM) - a single molecule localisation microscopy method allowing to determine the position of individual fluorescent molecules with nanometre accuracy in live cells (30). In PALM, tiny fractions of PvdA-PAmCherry are photo-activated at each acquisition frame, allowing determining their localisations with sub-diffraction precision. Thousands of sparse subsets of molecule coordinates pinpointed sequentially in time can then be accumulated within an imaged area. Using TIRF illumination limiting the observation the cytoplasmic zone immediately above the membrane contacting the surface, PvdA-PAmCherry was found to localise homogeneously in the cytoplasm. Despite a retrieved density of 2,000-3,000 molecule localisations per µm^2^, high enough to define spatial arrangements at a resolution of ∼40 nm resolution, no particular structural organisation was observed for PvdA, with the exception of preferential accumulation at one pole in some cells (25). We also used PALM to perform high-density single-particle tracking (sptPALM) to explore PvdA molecular trajectories within the cell. About 6,000 individual trajectories were computed from ∼54, 000 linkable individual localisations in 42 cells, allowing the construction of single molecule diffusion maps at the single cell level. Diffusion patterns appeared very similar from cell to cell (Figure 2B), with traces of heterogeneous velocities scattered throughout the cytoplasm (Figure 2C). We were not able to identify exclusion areas. To take advantage of the large number of short trajectories obtained with PvdA-PAmCherry in live cells, we computed jump-distances (JD) defined as the distance travelled by a single molecule during a fixed time lag Δ*t* (from 16 to 50 ms). Over the population of molecules, jump distance distribution reflects the fine features of the underlying diffusion mechanisms and/or the number of diffusing species (Figure 2 D)(31). JD histograms of PvdA were found very similar for the different cells. Assuming PvdA underwent Brownian motions, a nice adjustment of the empirical cumulative distribution of the JD was obtained with a two-populations model (Figure 2E), suggesting the presence of two diffusing populations of PvdA. Diffusion coefficients were 0.06 [0.02; 0.08] µm^2^ s^*-*1^ and 0.47 [0.39 – 0.54] µm^2^ s^*-*1^. We assigned these values to trapped and diffusing species, respectively (Figure 2 F). Due to the finite localisation precision (localisation error of ~ 30-40 nm), PvdA molecules diffusing at 0.06 µm^2^ s^*-*1^ could indiscriminately correspond to trapped, restrained or immobile fractions and were therefore referred as restrained hereafter. This fraction was found to represent about 10% to 24% (median 15%) of the total tracked PvdA. The value of 0.47 [0.39 – 0.54] µms^*-*2^ is about one order of magnitude slower as compared to the observed or calculated diffusion of free cytoplasmic proteins with the same molecular weight in *E. coli* (32). Although the viscosity of the cytoplasm of PAO1 might be slightly increased due to its smaller volume and its larger genome size as compared to *E. coli*, it is unlikely to explain such a difference. PvdA must therefore diffuse in the cytoplasm bound to a large complex.

**Fig. 2.**
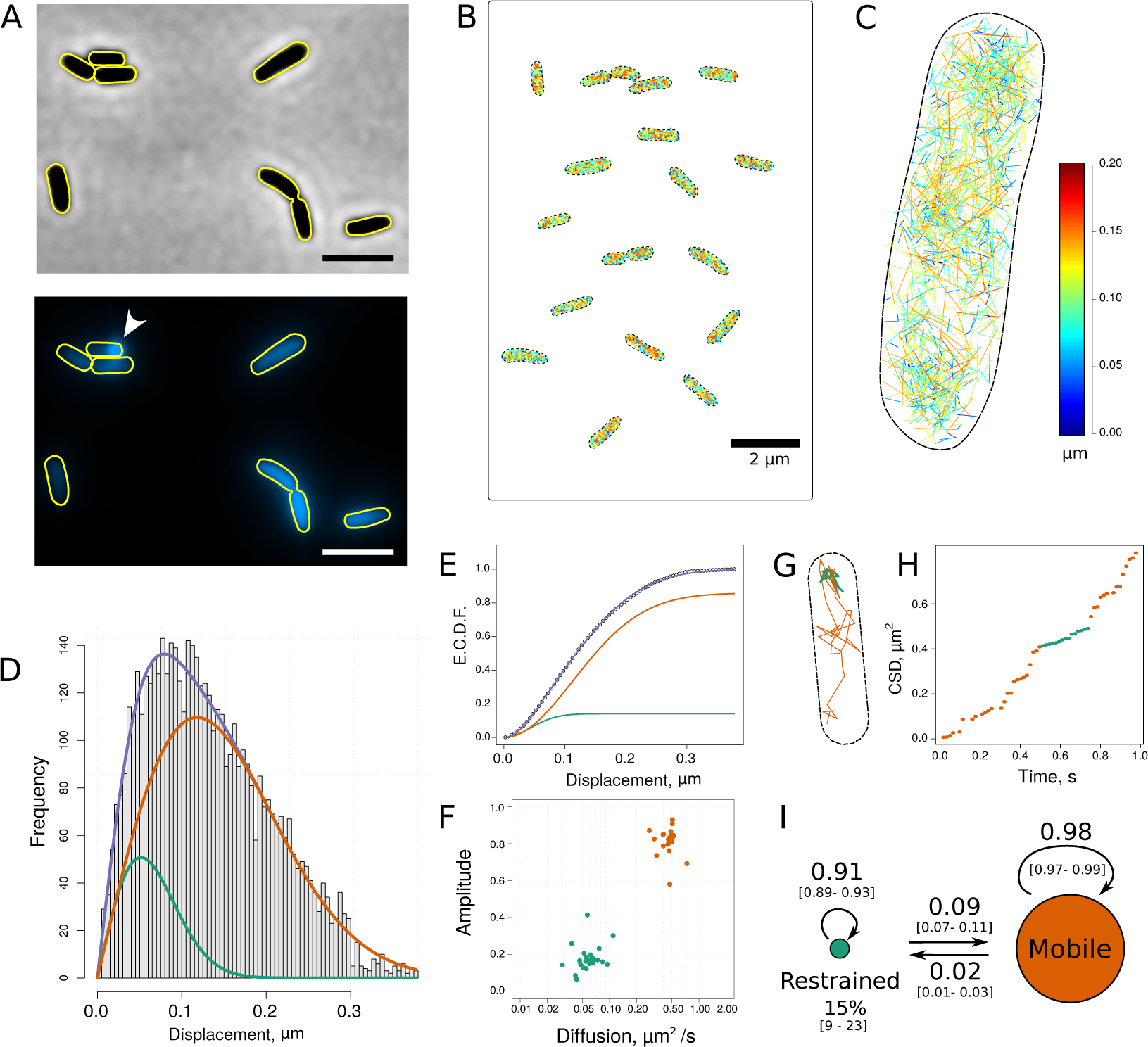
Single molecule tracking of PvdA-PAmCherry in live *P. aeruginosa*. (A) Phase-contrast (upper) and fluorescence (lower) images of PvdA-eGFP in PAO1 grown in Succinate Media at 30°C for 48 h. PvdA-eGFP is found homogeneously distributed in the cytoplasm. Some cells presented fluorescent spots at one pole (white arrow), as previously described (25–27). Scale bars = 2 µm. (B) Concatenated image of representative PvdA-PAmCherry cell diffusion maps obtained by sptPALM at 20°C and (C) magnification of a representative single cell diffusion map. Trajectories steps are colour-coded to indicate instantaneous jump distances (µm) observed during Δ*t* = 16 ms time interval. (D) Jump-distance distribution (JD) representing the Euclidean distance travelled by ∼ 23, 800 PvdA-PAmCherry molecules during a 16 ms time interval. These data correspond to the JD observed in 29 cells measured in four independent experiments. (E) The corresponding empirical cumulative distribution function (ecdf) was fitted assuming a two-population diffusion model. Diffusion coefficients of 0.06 [0.02 – 0.08] µm^2^ s^*-*1^ and 0.47 [0.39 – 0.54] µm^2^ s^*-*1^ (median [IQR]) were determined for the restrained (green) and mobile (orange) complexes, respectively. (F) Scatter plot of the amplitude of the restrained and mobile forms of PvdA-PAmCherry as a function of their corresponding diffusion coefficients (n = 23 cells). (G) Single PvdA-PAmCherry tracking trace observed at 62.5 Hz and (H) its cumulative square distance (CSD) as a function of time. The trace and the CSD are colour-encoded according to their diffusion regime defined by a discrete two-state Hidden Markov Model (HMM). (I) Schematic representation of the probability transition matrix retrieved from the HMM. The diameters of the diffusion regimen are proportional to their steady-state amplitudes. The probability of transition from mobile to restrained regimen was 2%. The bootstrapped 2.5% and 97.5% percentiles of the transition matrix parameters are in squared brackets.

To explore possible transitions between these two-state populations, we adjusted the square of the displacement values with a discrete two-state Hidden Markov models (HMM) assuming an exponential probability distribution. The HMM provides information about the average duration time in a state before transition, the probabilities for molecules to transition from one state to the other, and the mean diffusion value of each state. A representative tracking trace (although much longer than the 128 ms median trace duration) is presented in Figure 2G for illustration, together with its cumulative square displacement (CSD) plotted over time (32) 2H to provide a visual representation of the HMM model output. An overall linear increase of CSD for a given time window indicates that the jump-distances are distributed around their average value and therefore that the molecule diffuses uniformly. Break points defined by changes in slopes are likely to correspond to state transitions. It is clearly illustrated in figures 2G and H that the HMM model properly identifies the two states and their transitions. The transition probability matrix Pi of the model is presented in 2I. A PvdA in the restrained state has a probability of about 0.09 to transition to the mobile fraction on the next frame whereas only about 2% of the mobile PvdA are expected to become restrained. Although the probabilities are low, about 260 events of mobile-to-restrained and restrained-to-mobile transitions were observed within traces showing the two diffusion regimen are exchangeable. Finally, the amplitudes of the two populations were calculated as the steady-state probability distribution of the stationary Markov chain. A value of 15% (*CI*_95%_ = [9; 23]%) was associated to the constrained state, in very good agreement with the estimation made by fitting the JD empirical cumulative distribution. All these results were reproduced using PvdA-eYFP to rule out any quantification artefacts possibly resulting from PAmCherry photophysical properties (see supplementary Figure S2).

Taken together, these data clearly demonstrate that PvdA was mostly diffusing in the cytoplasm bound to large protein complexes without evidence for any spatial organisation pattern nor cytoplasmic exclusion areas at a ∼ 40 nm resolution (TIRF illumination). A small fraction estimated at ∼15% exhibited a reduced diffusion rate. We assigned it to a trapped complexes. The diffusing and trapped complexes were found exchangeable.

### PvdI interacts with many PvdA

PvdA is an ornithine hydroxylase (24) working in tandem with the hydroxy-ornithine transformylase PvdF (18) to generate fOH-Ornithine used as a building block by PvdI and PvdJ. To explore whether NRPS are cytoplasmic partners of PvdA, we sought to explore the physical interactions between PvdA and PvdI in live cells producing PVD. We used fluorescence lifetime imaging microscopy (FLIM) to measure Förster resonance energy transfer (FRET). FLIM is a general technique to probe changes in fluorophores’ local environment. In time-domain FLIM, a pulsed laser periodically excites the fluorophores in the sample. Following excitation, fluorophores relax into their ground-state either by emitting a photon (radiative decay) or through a non-radiative pathway. The fluorescence lifetime *τ* is defined as the average time a fluorophore remains in the excited state after excitation according to:

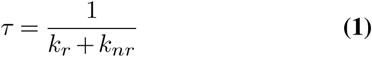

in which *k*_*r*_ and *k*_*nr*_ are the radiative and the non-radiative rate constants, respectively. Experimentally, each detected fluorescence photon is tagged with its arrival time relative to the last excitation laser pulse. The fluorescence lifetime *τ* can thus be retrieved by adjusting the experimental fluorescence decay with a sum of exponentials defined by rates inversely proportional to their fluorescence lifetimes. In the presence of a fluorescence acceptor (as mCherry) in the close vicinity of a fluorescence donor (as eGFP), FRET can occur-increasing the non-radiative de-excitation rate *k*_*nr*_ of the donor and thus decreasing its fluorescence lifetime. FRET can only occur if the acceptor is only few nm apart from the donor, a distance range implying physical interaction between the labelled molecules. FLIM FRET is therefore particularly well suited to probe protein-protein interactions in living cells (29, 33, 34). Moreover, as the fluorescence life-time is an intrinsic parameter of the fluorophore independent of concentration, FRET-FLIM is more robust than intensity-based methods to measure FRET when concentrations of interacting partners cannot be controlled.

Strains expressing only PvdA-eGFP or PvdI-eGFP cultured in succinate media for 48h were imaged using FLIM (Figure 3A). Their fluorescence decays were adjusted with a single exponential model to retrieve their average fluorescence life-time. The expression level of PvdA-eGFP, measured by the fluorescence intensity of cells on the microscopy images, was higher than that of PvdI-eGFP (see also supplementary Figure S3). But the fluorescence lifetime values of PvdA-eGFP or PvdI-eGFP were very similar with a median value of about 2.25 ns (Figure 3 B-C).

**Fig. 3.**
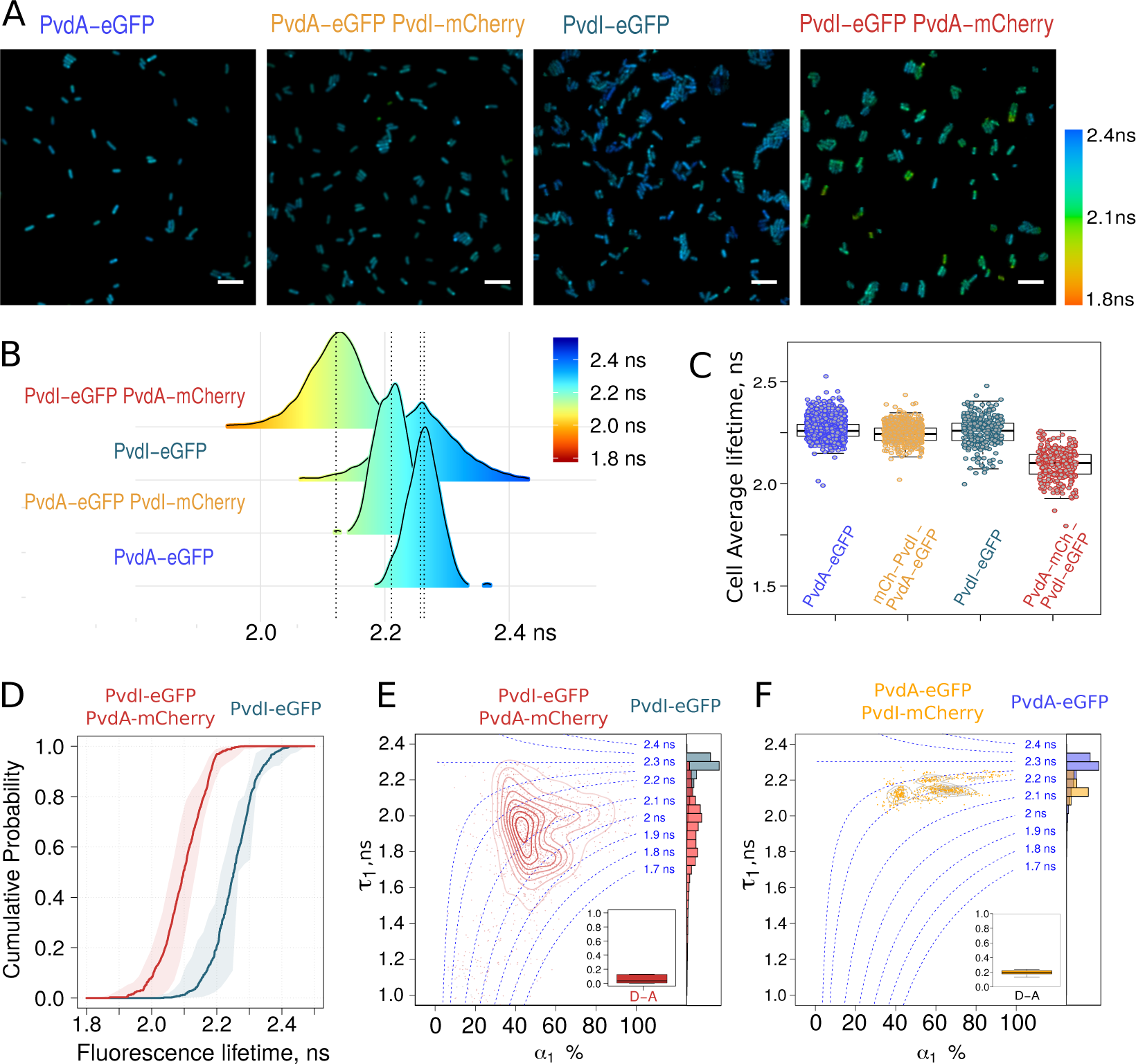
FRET-FLIM measurement of the PvdA-PvdI interaction in live PAO1. (A) Representative FLIM images of (from left to right) PvdA-eGFP, PvdA-eGFP PvdI-mCherry, PvdI-eGFP and PvdI-eGFP PvdA-mCherry strains, showing limited shortening in the average lifetime of PvdA-eGFP co-expressed with PvdI–mCherry in PAO1 but a larger effect for PvdI-eGFP co-expressed with PvdA-mCherry. All images are colour-encoded according to the pixel fluorescence lifetime values. The same colour scale (on the right) is used for the four images. Scale bar = 5 µm. (B) Probability density distribution of the mean fluorescence lifetime of pixels of (from bottom to top) PvdA-eGFP (n = 10,863), PvdA-eGFP PvdI-mCherry (n = 13,127), PvdI-eGFP (n = 6,962) and PvdI-eGFP PvdA-mCherry (n = 14,456) strains from at least N=3 independent experiments. (n = total number of pixels). (C) Average fluorescence lifetime of individual cells of (from left to right) PvdA-eGFP (n_cell_ = 1,154; N=8), PvdA-eGFP PvdI-mCherry (n_cell_ = 486; N=3), PvdI-eGFP (n_cell_ = 291; N = 3) and PvdI-eGFP PvdA-mCherry (n_cell_ = 303; N = 3). (n_cell_ = number of analysed cells; N = number of independent experiments). (D) Empirical cumulative probability distribution of the fluorescence lifetime of PvdI-eGFP and PvdI-eGFP PvdA-mCherry (data from panel B) (A), (B), (C) and (D) provide evidence for a physical interaction between PvdA and PvdI. (E) and (F) Density map with contour lines of (*τ* _1_, *α* _1_) fitting parameters showing clusters of pixels with similar decay signature for (E) PvdI-eGFP PvdA-mCherry (red) and (F) PvdA-eGFP PvdI-mCherry (orange). The histograms of the short-lived lifetime (*τ* _1_) of the donor-acceptor couple and the average lifetime (*τ*) of the corresponding donor-only conditions are plotted as marginal histograms. Inset: box plot of the fraction per cell of pixels better fitted with a single exponential decay model - corresponding to decays with no energy transfer.

Co-expression of PvdI-mCherry with PvdA-eGFP resulted in a limited shortening of the average lifetime of PvdA-eGFP (Figure 3 B-C),suggesting poor interaction or long inter-dyes distances. In sharp contrast, the average lifetime of cells expressing PvdI-eGFP was significantly shortened in the presence of PvdA-mCherry. Energy transfer between PvdI-eGFP and PvdA-mCherry can be evidenced by comparing the life-time distributions of PAO1 expressing PvdI-eGFP vs PvdI-eGFP PvdA-mCherry (Figure 3 B) or by comparing their corresponding empirical cumulative distributions (Figure 3 D). FRET can also be evidenced by comparing the lifetimes averaged by cells (Figure 3 C). It seems unlikely that the mere exchange of eGFP for mCherry or mCherry for eGFP has such a dramatic effect on PvdA/PvdI interaction. Indeed, both strains retain the ability of producing PVD with similar efficiencies (although lower than that of the wild-type PAO1 (see supplementary Figure S1)). The apparent contradiction in the above findings can be reconciled if we consider the expression levels of the two proteins with an expression of PvdA-eGFP far higher than that of PvdI-eGFP (see supplementary Figure S3). It can then be envisioned that PvdA-eGFP used as a donor coexists as a minor fraction transferring energy to mCherry and a major fraction free of transfer, both forms exhibiting different fluorescence lifetimes. Because the time resolved fluorescence decay sums photons emitted by the different species, the resulting fluorescence decay of PvdA-eGFP – PvdI-mCherry will thus tend towards the decay of PvdA-eGFP as the relative amount of non-transferring PvdA-eGFP increases. In contrast, the fluorescence decay of cells co-expressing PvdI-eGFP and PvdA-mCherry will be shifted to lower lifetimes, as the less abundant PvdI-eGFP protein is transferring to the nearby PvdA-mCherry.

To take this species mixture into account, fluorescence decays were fitted with a two-exponential component model in which the average lifetime is defined as a weighted sum of two decays according to:

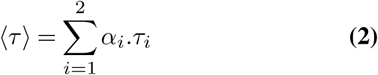

To limit over-fitting and improve fitting convergence, the long-lived lifetime *τ*_2_ was fixed at the 90^th^ percentile of PvdA-eGFP or PvdI-eGFP lifetimes defined using a single exponential decay (2.25 - 2.30 ns), whereas the lifetime value of the fast decay *τ*_1_ and the relative contribution of each component defined b y *α* _1_ a nd *α* _2_ could float. For a two-component model *α*_2_=1-*α*_1_ and ⟨*τ*⟩ can thus be rewritten as

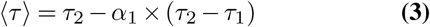

A successful fit procedure will converge toward ⟨*τ*⟩ by adjusting *τ*_1_ and *α*_1_ and will provide *τ*_1_ and *α*_1_ conditional probability distributions given the fixed *τ*_2_ value. A plot of *τ*_1_ as a function of *α*_1_ can be drawn as a diagram to represent observations made over all fitted pixels. An estimation of the 2D density of the pixel spatial distribution in this plot provides an estimation of the average lifetime (centre of the density map contour plot) but also the bivariate distribution of (*τ*_1_, *α*_1_) parameters. The distribution of (*τ*_1_, *α*_1_) measurements of PvdI-eGFP PvdA-mCherry was scattered with a significant part of the distribution showing shortened *τ*_1_ as compared to PvdI-eGFP lifetime - with values ranging from about 2.2 ns down to 1.6 ns (Figure 3 E). This confirmed a pronounced energy transfer between the two fluorophores. The fraction of pixel decays better adjusted with a single exponential was only about 5% (Figure 3 E inset). These FLIM data clearly showed that PvdA is physically interacting with PvdI.

In sharp contrast, the distribution of (*τ*_1_, *α*_1_) parameters of PvdA-eGFP – PvdI-mCherry was mostly positioned in an area corresponding to an average lifetime of about 2.2 ns with *τ*_1_ values about 2.1 ns (Figure 3 F). If equimolar complex of PvdA and PvdI were formed, the two (*τ*_1_, *α*_1_) maps for PvdI-eFGP – PvdA-mCherry and PvdA-eGFP – PvdI-mCherry should look very similar, only being shifted along the *α*_1_ axis due to the difference in the relative amount of energy transferring species. Indeed, if we consider the difference in the expression level of the two enzymes, almost all the PvdI molecules should be bound to PvdA (high value for *α*_1_) while only a small fraction of PvdA should be bound to PvdI (low value of *α*_1_) but in both cases *τ*_1_ should not be affected because the structure of the interacting complex should remain the same. Our observations largely differed from this situation with *τ*_1_ values being also affected. This difference is likely explained by the presence of multiple acceptors, further shortening the average lifetime according to:

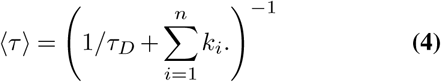

in which *n* is the number of acceptors and *k*_*i*_ and *τ*_*D*_, the rate of energy transfer and the donor lifetime, respectively. The short *τ*_1_ values observed for PvdI-eGFP PvdA-mCherry (compare red contour lines in Figure 3E and orange ones in Figure 3 F) are likely explained by the presence in the complexes of several PvdA-mCherry surrounding PvdI-eGFP in a range of distances compatible with FRET. One can thus infer that multiple PvdA-mCherry proteins are bound per PvdI-eGFP NRPS.

Taken together, these observations provide evidence for a physical interaction between PvdA and PvdI and suggest that multiple PvdA are bound to one molecule of PvdI.

### PvdA interacts with PvdJ

As PvdJ also contains a module responsible for the addition of L-fOH-Ornithine in the pyoverdine peptide, we decided to monitor FLIM-FRET between PvdA and PvdJ (Figure 4 A). The lifetime of PvdJ-eGFP PvdA-mCherry estimated from a single exponential decay model was shortened by about 10-15% (2.0-2.1 ns) as compared to PvdJ-eGFP fluorescence lifetime (Figure 4A). As for PvdA-PvdI interaction, we could not define any preferential area where the interaction takes place. Using a two-exponential decay model, the fraction of decays better adjusted with a single exponential model was close to zero (Figure 4 B inset). The distribution of *τ*_1_ was centred at about 2.0 ns and the amplitude of the short-lived component was representing about 60% to 80%. Interestingly, a similar *τ*_1_ value was observed for the fraction of PvdA-eGFP molecules interacting with PvdJ-mCherry but associated to a *α*_1_ value of ∼40%. Therefore, the same *τ*_1_ value observed for the both labelling schemes of the two enzymes. This strongly suggests the presence of a unique complex with a well-defined binding site for PvdA on PvdJ that governs the interaction. This conclusion is in line with a previous report showing by double-hybrid experiment that PvdA interacts with PvdJ M2 isolated module (26) - the module responsible for incorporation of formyl-OHOrn (Figure 1 C). The high value of *α*_1_ observed for PvdJ-eGFP PvdA-mCherry indicated that in the cellular context, the binding site of PvdJ is saturated by PvdA. Unfortunately, we cannot infer from our data if it corresponds to a site with strong affinity and a long residence time of PvdA on PvdJ or to a situation of fast exchange of PvdA molecules on this binding site.

**Fig. 4.**
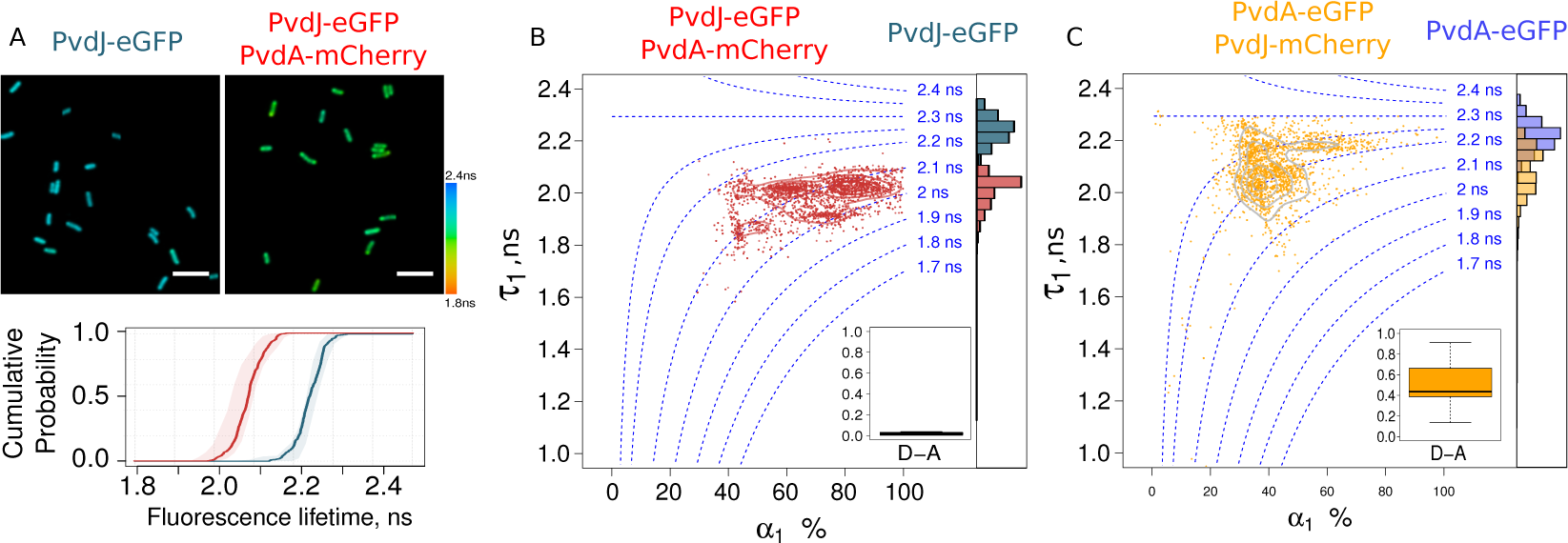
FRET-FLIM measurement of the PvdA-PvdJ interaction in live PAO1. (A) Representative FLIM images of PvdJ-eGFP and PvdJ-eGFP PvdA-mCherry strains. Using a single exponential analysis, a shortening of the lifetime of PvdJ-eGFP co-expressed with PvdA–mCherry is observed in live PAO1. The cumulative probability of the fluorescence lifetime of these two strains demonstrates the interaction of PvdA with PvdJ in live cells. (B) Density map with contour lines of (*τ* _1_, *α* _1_) fitting parameters of PvdJ-eGFP (grey) and PvdJ-eGFP PvdA-mCherry (red). (C) Density map with contour lines of (*τ* _1_, *α* _1_) fitting parameters of PvdA-eGFP (blue) and PvdA-eGFP PvdJ-mCherry (orange). (B) and (C) provide evidence for a major and possibly unique binding site for PvdA on PvdJ

**Fig. 5.**
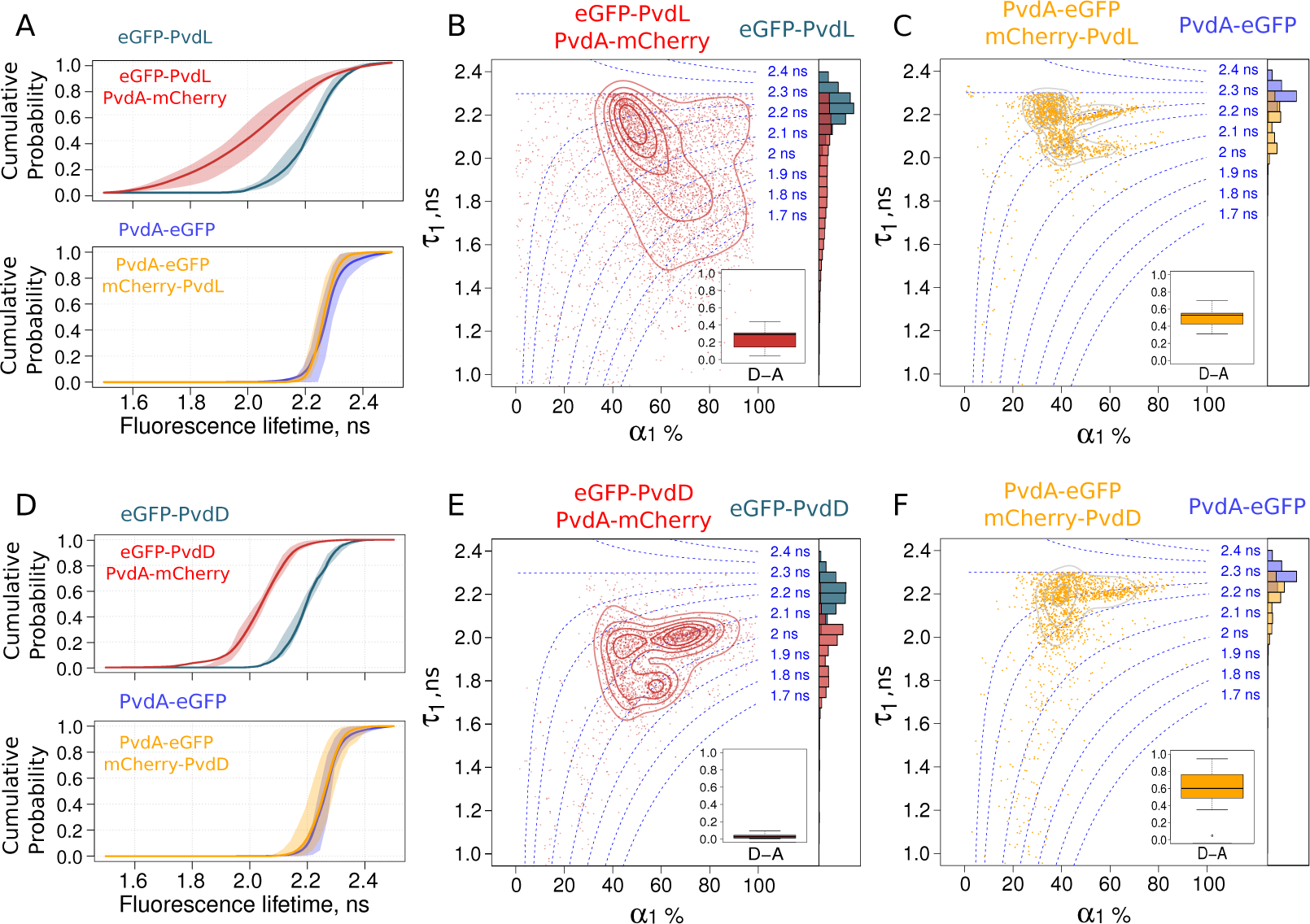
FRET-FLIM measurement of the PvdA-PvdL and PvdA-PvdD interactions in live PAO1. Cumulative probability of the fluorescence lifetime of the PvdA-PVdL (A) and PvdA-PvdD complexes (D). Larger fluorescence lifetime decrease were observed when PvdL or PVdD were labelled with eGFP. Density map with contour lines of (*τ* _1_, *α* _1_) fitting parameters of (in B) eGFP-PvdL (grey) and eGFP-PvdL PvdA-mCherry (red) and (in C) PvdA-eGFP (blue) and PvdA-eGFP mCherry-PvdL (orange). Density map with contour lines of (*τ* _1_, *α* _1_) fitting parameters of (in B) eGFP-PvdD (grey) and eGFP-PvdD PvdA-mCherry (red) and (in C) PvdA-eGFP (blue) and PvdA-eGFP mCherry-PvdD (orange).

### PvdA interacts with PvdL and PvdD in cells

We further investigated the interaction of PvdA with PvdL (5A-C) and PvdD (5 D-F). None of these two NRPS are using PvdA product as a building block. We were nevertheless able to evidence physical interactions between PvdA and PvdL (5A-C) and between PvdA and PvdD (5D-E). It clearly appeared that the binding mode of PvdA to PvdL differed from the binding of PvdA to PvdD. It was also visible that results observed for PvdL (5A-C) were very similar to that observed for PvdI (3D-F). On the other hand, observations made for PvdD (5D-F) were broadly like PvdJ (4A-C) with a significant amount of interacting eGFP-PvdD characterised by a *τ*_1_ value of about 2.0 ns – the same *τ*_1_ value than that measured for the small fraction of PvdA-eGFP interacting with mCherry-PvdD 5 panel F. But in contrast to PvdJ, a second binding site with shorter *τ*_1_ value (about 1.8 ns) was also present (5E)– suggesting that PvdD can accommodate more than one PvdA protein.

## Discussion

In this work, we constructed PAO1 strains with modified genes coding for PvdA and NRPS fluorescent proteins (see supplementary table S1) inserted at the chromosomal level. These tagged enzymes retained the expression levels and functions of the endogenous ones. We used these strains to explore interactions between PvdA and the four NRPS responsible for the bio-synthesis of the PVD peptide backbone. We found that PvdA was not existing as a free protein in the cytoplasm but was engaged in large complexes. We inferred from FLIM data the physical interactions of PvdA with PvdI and PvdJ but also with PvdL and PvdD and demonstrated different binding modes of PvdA on the different NRPS. Based on the differences in FRET distributions between strains expressing PvdA tagged with the fluorescent donor (PvdA-eGFP) as compared to strains expressing PvdA tagged with the acceptor mCherry, we evidenced the simultaneous binding of multiple PvdA bound to several NRPS and clearly distinguished this situation from complexes with 1:1 stoichiometry. Interestingly, the interaction of PvdA with PvdI and with PvdJ, the two NRPS using fOHOrn as a building block, differed in extent and nature. The presence of several binding sites has to be considered to explain our FLIM observations with PvdI. The interaction of several PvdA with PvdI may fulfil the need for an excess of substrate locally available to optimise the activity of the PvdI enzyme. An unbalanced binding stoichiometry may also compensate fOHOrn dilution in the cell cytoplasm. On the contrary, a major binding site for PvdA was found on PvdJ. The M2 module previously identified by two-hybrid as an interacting partner of PvdA (26) and responsible for fOHOrn insertion is a good candidate to harbour this binding site for PvdA. The difference of binding mechanism of the two NRPS is striking if we consider that both PvdI and PvdJ require the action of PvdA. Since PvdL exhibits a global binding behaviour very similar to PvdI, the molecular size of the NRPS could be an element explaining the binding mode – the larger the NRPS the higher the number of PvdA binding sites the enzyme can accommodate. Additional speculations can be made. For instance, a connection could be made between (i) the conserved position of fOH-Orn in the PVD sequence of *Pseudomonas aeruginosa* and (ii) the presence or not of a single or multiple binding sites. Pyoverdine production is a characteristic of the fluorescent pseudomonads. But *Pseudomonas* species produce slightly different pyoverdines. *P. aeruginosa* strains are able to produce three different pyoverdines, PVD I, PVD II and PVD III with a different peptide sequences. Amongst the pyoverdine sequences of *P. aeruginosa*, the module 2 of PvdJ is well conserved and the fOHOrn is highly conserved at that position 9 in the PVD precursor sequence. On the contrary, the position where the fOHOrn is inserted by PvdI in the PVD precursor is more variable and corresponds either to module 2, 3 or 4 of PvdI in the different *P. aeruginosa* strains (7, 35). In this context, evolving multiple binding sites outside the modules provide the necessary flexibility to ensure that the product produced by PvdA is available whatever the fOH ornithine module position within PvdI. It suggests additionally that rather than physical coordination between active sites of tailoring enzymes and NRPS modules, only co-localisation of sequential enzymes seems to be enough to promote metabolic efficiency. Taken together, we can then envisage binding sites have co-evolved with module specificity.

Although we did not interrogate direct interactions between the different NRPS in this work, our findings provide strong evidence for siderosomes. Such organisation has been previously hypothesised based on biochemical observations, but the difficulty to handle these complexes limits their *in vitro* explorations. The reversibility of spatial assemblies of sequential metabolic enzymes - proposed as a general mechanism to regulate metabolic activities-may explain why their *in-vitro* study remains extremely challenging. Our data demonstrate the possibility to capture rich information on protein-protein interactions which allows to directly characterise siderosomes in the cellular environment even if their formation is transient. This type of observation is susceptible to provide valuable data regarding the importance of maintaining effective protein–protein interactions in recombinant NRPS templates, or more generally recombinant metabolic machinery. But our work also raised some questions in particular regarding the slow diffusion of PvdA and the fraction with restrained diffusion. Does it correspond to complexes interacting with PvdE, the transporter responsible for the PVD precursor export to the periplasm? Are siderosome interacting with the inner membrane and therefore can they be considered as metabolons? In both cases, the possible additional spatial regulation resulting from specific partitioning within the bacterial cytoplasm will require 3-dimensional multi-colour explorations in the cellular context.

Taken together, our data clearly evidence that secondary metabolite pathways are much more organised than initially envisioned, with multiple interactions and multiple binding modes between accessory proteins and NRPS. If clustering of all the PVD bio-synthesis machinery in a single complex is possibly increasing the synthesis efficiency, notably by sequestrating intermediate products, the understanding of how metabolic enzymes interact and coordinate with each other in space and time to ensure their metabolic functions is mostly at its infancy. But due to their non-vital nature and their experimentally tractable expression, siderosomes are good candidates to be used as model systems to demonstrate the functional significance of highly orchestrated enzyme organisations in the cellular context. Additional general knowledge gained from siderosome will likely contribute further to the general understanding of mechanisms ruling widespread metabolic pathways and cellular organisations.

## Material and Methods

### Bacterial strains, plasmids and growth conditions

*P. aeruginosa* strains used in this study are listed in supplementary Table S1. Bacteria were grown in 5 mL lysogeny broth (LB) (Difco) at 30°C under 220 rpm orbital shaking. To induce the expression of the pyoverdine pathway enzymes, *P. aeruginosa* strains were pelleted by centrifugation, washed and further grown overnight at 30°C in an iron-deficient succinate medium (composition: 6 g L^*-*1^ K_2_HPO_4_, 3 g L^*-*1^ KH_2_PO_4_, 1 g L^*-*1^ (NH_4_)_2_ SO_4_, 0.2 g L^*-*1^ MgSO_4_, 7 H_2_O and 4 g L^*-*1^ sodium succinate with the pH adjusted to 7.0 by adding NaOH). Cells were finally diluted and grown an additional 24 hours. The presence of pyoverdine in the supernatant can be visually observed as an intense yellow-green water-soluble pigment.

### Mutants construction

All enzymes for DNA manipulation were purchased from ThermoFisher Scientific and were used in accordance with the manufacturer’s instructions. *E. coli* strain TOP10 (Invitrogen) was used as the host strain for all plasmids. The DNA fragments from *P. aeruginosa* used for cloning were amplified from the genomic DNA of the PAO1 strain with Phusion High-Fidelity DNA polymerase (ThermoFisher Scientific). The primers used are listed in supplementary Table S3 of the Supporting Information. As previously described (25), the general procedure for the construction of the plasmids involved insertion of the *mcherry, egfp* or *pamcherry* gene flanked by upstream and downstream regions of 700 bp, corresponding to the insertion site, into a pME3088 or a pEXG2 vector. Mutations in the chromosomal genome of *P. aeruginosa* were generated by transferring the suicide vector from *E. coli* TOP10 strain into the *P. aeruginosa* recipient strain and allowing the plasmid to integrate into the chromosome, with selection for tetracycline resistance for pME3088 (36) or gentamicin resistance for pEXG2. A second crossing-over event excising the vector was achieved by enrichment for tetracycline-sensitive cells (pME3088) or sucrose resistant cells selection (pEXG2), to generate the corresponding mutants (37). All tag insertion mutants were verified by PCR and sequencing.

### Wide field imaging

Wide field imaging was performed on a Micro-Manager (38) controlled Olympus IX-81 inverted microscope equipped with z-drift control and auto-focus system (ZDC Olympus). Phase images were generated using a Phase3 100X 1.4NA objective (Olympus) using a 512 × 512 pixels EM-CCD (ImagEM - Hamamatsu Photonics - Japan). Epifluorescence excitation was provided by a 488 nm laser diode (spectra physics) using a 488 nm (Di01-R488 - Semrock) dichroic filter. Fluorescence signals were filtered using a 488 nm Long pass filter (BLP01-488R-25 – Semrock). Live-cells were immobilised on a 1% agarose pad and imaged at 20 °C. Quantitative fluorescence measurements were performed at the cell level using a home-made imageJ plugin for cell segmentation and fluorescence quantification. Initial cell segmentation was based on a Laplacian of Gaussian (LoG) filtering of the phase image and further used to define a rod-shape contour (sphero-cylinder) based on different central moment calculations. Initial rod-shape contours were then refined based on the isotropic 3 × 3 gradient of the initial image to adjust bacteria contours.

### Photo-Activated Localisation Microscopy (PALM) and single particle tracking-PALM (sptPALM)

PALM and sptPALM (28) were performed on a home-built Micro-Manager (38) controlled inverted-microscope based on a Nikon Eclipse II equipped with a 100X 1.49 objective (Nikon - Japan) and a drift focus compensator (Perfect Focus System - Nikon-Japan). Photo-activable mCherry (PA-mCherry) was excited with a 561 nm diode laser (Oxxius-France) and activated with a 405 nm laser diode (Spectra Physics - Germany) selected by an acousto-optic tunable filter (AOTFnC 400-650 TN, AA Otpo Electronic-France). Fluorescence emission of PA-mCherry was collected through a quadri-band dichroic filter (FF405/496/560/651-Di01-25×36-Semrock - USA) and further filtered with a 561 nm long-pass filter (561nm RazorEdge, Semrock - France). The wavefront distortions of the emission signal were corrected using an adaptive optics device (MicAO, Imaging Optics - France) before being imaged on a 512 × 512 pixels EM-CCD (ImagEM - Hamamatsu Photonics - Japan) operating at - 65 °C with an ADU to photon conversion factor of about 60. Image stacks ranging from 2000 to 5000 frames with an exposure time of 16 to 50 ms were typically recorded for sptPALM using a home-written Beanshell acquisition script. Molecules localisations were retrieved using Peakfit, an open source software package for Fiji and further analysed using self-written script in R. Tracking traces were generated using the simple Linear Assignment Problem (LAP) tracker (39) implemented in TrackMate plugin (40) for Fiji. The simple LAP is well suited for tracking molecules undergoing Brownian motion or for low density of molecules undergoing non-Brownian motion. Data were further analysed in R according to a jump distance analysis (31) assuming the mean square displacement is proportional to the two-dimensional diffusion coefficient D (in µm^2^*/*s) (2-D Brownian motion) such that ⟨*r*^2^(Δ*t*)⟩ = 4*D*_*j*_Δ*t*. Diffusion coefficients were retrieved from the empirical cumulative distribution function (ecdf) of the particles’ displacements in a set time interval according to :

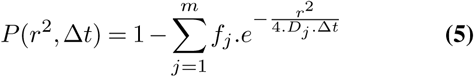

The choice the ecdf instead of the discrete probability density function (pdf) histogram for fitting was motivated by the dependence of the pdf histogram to the user input for bin-size selection - possibly biasing the fitting parameters. Hidden Markov model estimates were obtained using the Baum-Welch algorithm (41) and the Viterbi algorithm (42), both implemented in the R hmm package (https://cran.r-project.org/package=HMM) that predict the most likely sequence of Markov states given the observed dataset. Diffusion coefficients were obtained from the HMM exponential rates. HMM parameter confidence intervals (exponential rates and transition matrix) were obtained by bootstrap.

The tracking analysis procedure was challenged using data of the Particle Tracking Challenge (http://www.bioimageanalysis.org/track/) found in Supplementary Video 1 of reference (43) corresponding to simulated images of vesicles undergoing Brownian motions with a medium particle density and a signal-to-noise ratio of four. Retrieved parameters were in excellent agreement with the ground truth as shown in Supplementary Figure S5.

### Fluorescence Lifetime Imaging Microscopy (FLIM)

Time-correlated single-photon counting FLIM measurements were performed on a home-made two-photon excitation scanning microscope based on an Olympus IX70 inverted microscope with an Olympus 60× 1.2NA water immersion objective operating in de-scanned fluorescence collection mode (44, 45). Two-photon excitation at 930 nm was provided by a Ti:Sapphire oscillator (Mai Tai® DeepSee™, Spectra Physics - 80 MHz repetition rate, *≈* 70 fs pulse width) at 10-20mW. Fluorescence photons were collected through a 680 nm short pass filter (F75-680, AHF, Germany) and a 525/50 nm band-pass filter (F37-516, AHF, Germany) and directed to a fibre-coupled avalanche photo-diode (SPCM-AQR-14-FC, Perkin Elmer) connected to a time-correlated single photon counting (TCSPC) module (SPC830, Becker & Hickl, Germany). Cells grown for 48h in succinate media were immobilised on a 1% agarose pad and rapidly imaged. Typically, area of 50×50 µm in the samples were scanned at 4 µs per pixel (1024 × 1024 pixels) for 100 s to 600 s to reach the Nyquist-Shannon sampling criteria and to achieve the appropriate photon statistics to investigate the fluorescence decays. Fluorescence decays were processed using a commercial software package (SPCImage V2.8, Becker & Hickl, Germany). A binning of two was applied before processing the fluorescence decays. FLIM data were further analysed using a homemade ImageJ plugin and R scripts.

## ACKNOWLEDGEMENTS

We acknowledge Dr Ludovic Richert for his valuable assistance on FLIM data acquisition and for the technical maintenance and development of the FLIM setup. YM is grateful to the Institut Universitaire de France (IUF) for support and providing additional time to be dedicated to research.

