## Supplementary tables and figures for "Supra-molecular organization of the pyoverdine bio-synthetic pathway in *Pseudomonas aeruginosa*"

**Supplementary materials**

Table S1: *P. aeruginosa* strains used in this study.

| Strains | Collection ID | Relevant characteristics | Source |
|---|---|---|---|
| *Pseudomonas aeruginosa* | | | |
| PAO1 | **PAO1** | Wild-type strain | Stover *et al.*[1] |
| PvdA-mCherry | **PAS159** | Derived from PAO1 - chromosomally integrated | Gasser et al.[2] |
| PvdI-mCherry | **PAS178** | Derived from PAO1 - chromosomally integrated | This work |
| PvdA-eGFP | **PAS180** | Derived from PAO1 - chromosomally integrated | Gasser et al.[2] |
| PvdA-eGFP mCherry-PvdL | **PAS181** | Derived from PAO1 - chromosomally integrated | Gasser et al.[2] |
| PvdA-eGFP mCherry-PvdD | **PAS186** | Derived from PAO1 - chromosomally integrated | Gasser et al.[2] |
| eGFP-PvdD | **PAS214** | Derived from PAO1 - chromosomally integrated | This work |
| eGFP-PvdL | **PAS215** | Derived from PAO1 - chromosomally integrated | This work |
| PvdI-eGFP | **PAS216** | Derived from PAO1 - chromosomally integrated | This work |
| eGFP-PvdD PvdA-mCherry | **PAS229** | Derived from PAO1 - chromosomally integrated | This work |
| eGFP-PvdL PvdA-mCherry | **PAS230** | Derived from PAO1 - chromosomally integrated | This work |
| PvdI- eGFP PvdA-mCherry | **PAS231** | Derived from PAO1 - chromosomally integrated | This work |
| PvdA-eGFP mCherry-PvdI | **PAS246** | Derived from PAO1 - chromosomally integrated | Gasser et al.[2] |
| PvdA-eGFP PvdJ-mCherry | **PAS247** | Derived from PAO1 - chromosomally integrated | Gasser et al.[2] |
| PvdA-PAmCherry | **PAS405** | Derived from PAO1 - chromosomally integrated | This work |
| PvdA-eGFP PvdI-mCherry | **PAS446** | Derived from PAO1 - chromosomally integrated | This work |
| PvdJ-eGFP | **PAS471** | Derived from PAO1 - chromosomally integrated | This work |
| PvdJ-eGFP PvdA-mCherry | **PAS472** | Derived from PAO1 - chromosomally integrated | This work |
| *Escherichia coli* | | | |
| TOP10 | | *F- mcrA Δ(mrr-hsdRMS-mcrBC) φ80lacZΔM15 ΔlacX74 nupG recA1 araD139 Δ(ara-leu)7697 galE15 galK16 rpsL*($Str^R$) *endA1* $\lambda^-$ | Invitrogen |

Table S2: Plasmids used in this study.

| **Plasmids** | **Collection ID** | **Relevant characteristics** | **Source** |
|---|---|---|---|
| pEXG2 PvdA-PAmCherry | **pAF10** | pEXG2 carrying the sequence to insert a PA-mCherry tag in Cter of *pvdA* | This work |
| pEXG2 | **pEXG2** | Allelic exchange vector with pBR origin, gentamicin resistance, *sacB* | Rietsch *et al.*[3] |
| pME3088 | **pME3088** | Allelic exchange vector with ColE1 origin, tetracyclin resistance | Voisard *et al.*[4] |
| pME3088 PvdI-mCherry | **pLG42** | pME3088 carrying the sequence to insert a mCherry tag in Cter of *pvdI* | This work |
| pME3088 eGFP-PvdD | **pVEGA15** | pME3088 carrying the sequence to insert a eGFP tag in Nter of *pvdD* | This work |
| pME3088 eGFP-PvdL | **pVEGA16** | pME3088 carrying the sequence to insert a eGFP tag in Nter of *pvdL* | This work |
| pME3088 PvdI-eGFP | **pVEGA17** | pME3088 carrying the sequence to insert a eGFP tag in Cter of *pvdI* | This work |
| pEXG2 PvdJ-eGFP | **pVEGA30** | pEXG2 carrying the sequence to insert a eGFP tag in Cter of *pvdJ* | This work |
| pEXG2 PvdA-eGFP | **pAF8** | pEXG2 carrying the sequence to insert a eGFP tag in Cter of *pvdA* | This work |
| pEXG2 PvdA-PAmCherry | **pAF10** | pEXG2 carrying the sequence to insert a PA-mCherry tag in Cter of *pvdA* | This work |

Table S3 Primers used in this study.

| Oligonucleotides | Sequence (5' to 3') | Used to construct the following plasmids |
|---|---|---|
| PvdI-XhoIFC | AAACTCGAGTTCGTGCCGGATCCCTTTG | pLG42 |
| PvdI-XbaIRC | TTTTCTAGAGATCGCCTCTAGTTCGCTC | pLG42 |
| PvdI-ClaIFC | AAAATCGATTGACCCATGCTTTCCAATCCA | pLG42 |
| PvdI-HindIIIRC | TTTAAGCTTGCCGGTCCAGTACGCCAACTG | pLG42 |
| mCHE-XBAF | AAATCTAGAGTGAGCAAGGGCGAGGAG | pLG42, pVEGA15, pVEGA16, pVEGA17 |
| mCHE-CLAR | AAAATCGATCTTGTACAGCTCGTCCAT | pLG42, pVEGA15, pVEGA16, pVEGA17 |
| PvdD-EcoRIFN | AAAGAATTCGGATGGGGTGGTGGACTACCTC | pVEGA15 |
| PvdD-XbaIRN | TTTTCTAGACACGCTACCGCCTCTTAGGAAATC | pVEGA15 |
| PvdD-ClaIFN | AAAATCGATCAAGCACTCATAGAGAAGGTG | pVEGA15 |
| PvdD-HindIIIRN | TTTAAGCTTGCCCAGCAGGCCGGTCCAG | pVEGA15 |
| PvdL-HindIIIFN | AAAAAGCTTTTCGGCGAGGCCCTGCATACCG | pVEGA16 |
| PvdL-XbaIRN | TTTTCTAGACATCATGTGTTTTCCTGCCTG | pVEGA16 |
| PvdL-ClaIFN | AAAATCGATGACGCCTTCGAACTTCCCACC | pVEGA16 |
| PvdL-XhoIRN | TTTCTCGAGTACGCCGCTGAAGATCGGTTG | pVEGA16 |
| PvdI-XhoIFC | AAACTCGAGTTCGTGCCGGATCCCTTTG | pVEGA17 |
| PvdI-XbaIRC | TTTTCTAGAGATCGCCTCTAGTTCGCTC | pVEGA17 |
| PvdI-ClaIFC | AAAATCGATTGACCCATGCTTTCCAATCCA | pVEGA17 |
| PvdI-HindIIIRC | TTTAAGCTTGCCGGTCCAGTACGCCAACTG | pVEGA17 |
| egfpF | gtgagcaagggcgaggagctgttcaccgggg | pVEGA30 |
| egfpR | cttgtacagctcgtccatgccgagagtgatcccgg | pVEGA30 |
| pvdJstop-832F | GTACCTGGGCGGGGAAGGGGTGGCGCGT | pVEGA30 |
| pvdJstop+852R | GGGCCACGCGCCTGAGCGCCTGGGAC | pVEGA30 |
| egfppvdJoverlapF | ccgggatcactctcggcatggacgagctgtacaagTAAGAGGCGGTAGCGTGCAAGCACTCATAGAGAAGGTGG | pVEGA30 |
| pvdJegfpoverlapR | ccccggtgaacagctcctcgcccttgctcacGGAAATCAGTTTTTCAAGTTCATCGGCAGATAGACGTTTGAGCGCCTC | pVEGA30 |
| PvdA stop-700 EcoRI For | ATCGGAATTCGATGAAGATCGCCATTATCGG | pAF8, pAF10 |
| PvdA stop-700 Rev | GCTGGCCAGGGCGTGCT | pAF8, pAF10 |
| eGFP For overlap PvdA | AGCACGCCCTGGCCAGCGTGAGCAAGGGCGAGGA | pAF8, pAF10 |
| eGFP Rev | CTTGTACAGCTCGTCCATGC | pAF8, pAF10 |
| PvdA Stop+700 For over GFP | GCATGGACGAGCTGTACAAGTGATCGGCGCCACGCCG | pAF8, pAF10 |
| PvdA Stop+700 Hind Rev | ATCGAAGCTTCAAGGCGACCTTCTCCGC | pAF8, pAF10 |

Figure S1

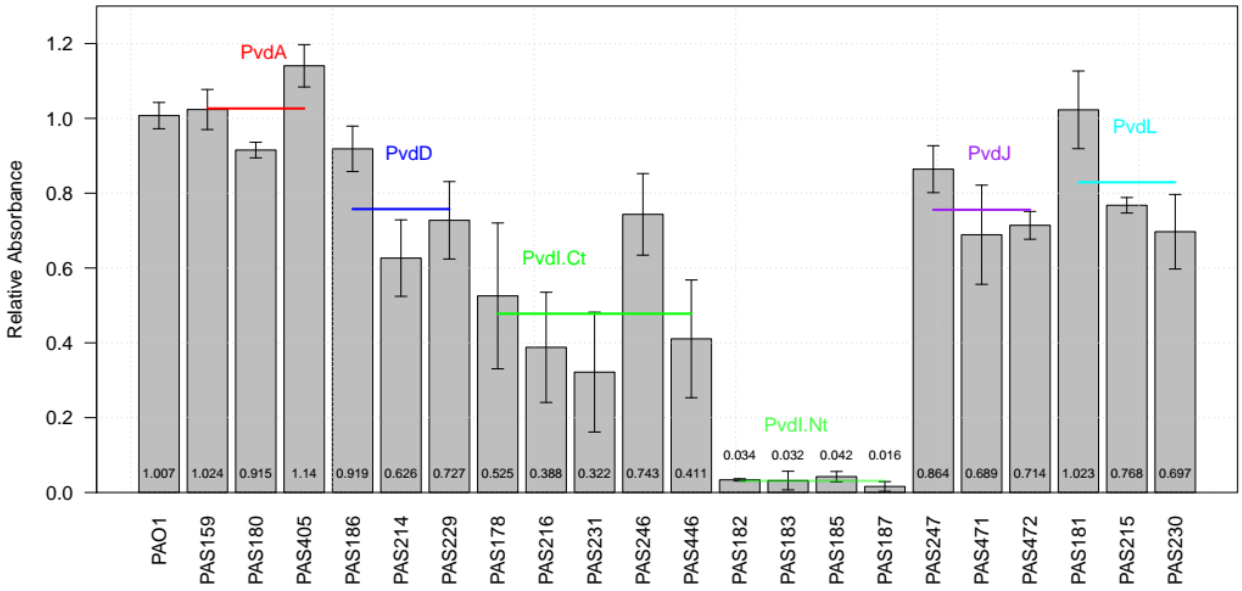


Figure S1

**PVD production of the different strains used in this study.**

PVD production was followed by measuring the corrected relative absorbance at 405 nm (compared to PAO1) of the filtered supernatant of the succinate growing media (SM). The absorbance was measured on cultures grown at 30°C for 48h in SM. Strains are organized on the graphic according to the labelled protein (see supplementary table S1 for details) – doubly-labelled strains (PvdA and NRPS labelling) were assigned to NRPS groups. Note that strains with PvdI modified at their N termini were not used in this study as these modifications were interfering with PVD production.

Figure S2

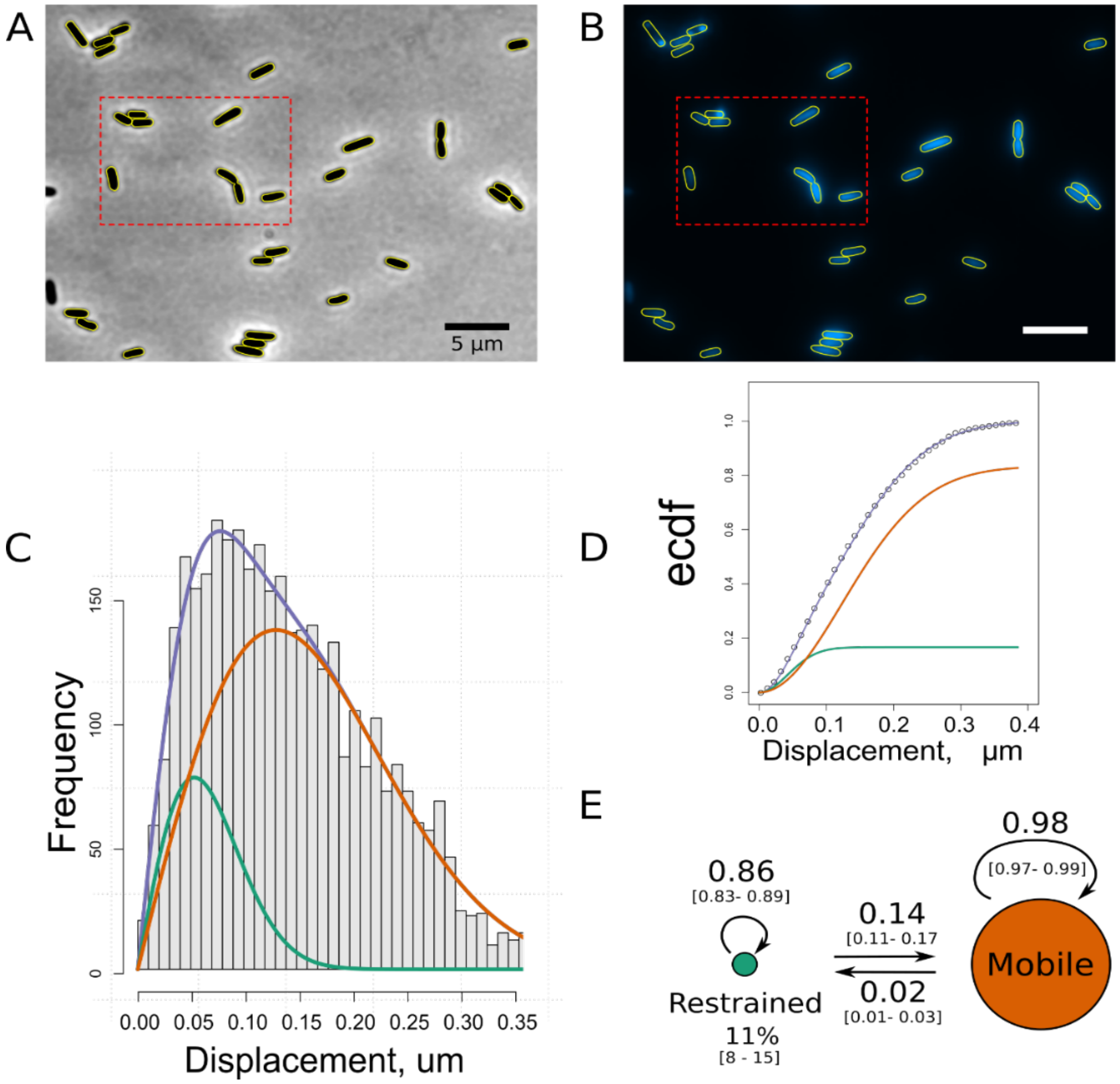


*Figure S2*

**Single molecule tracking of PvdA-eYFP in live *P. aeruginosa***

(A) Phase-contrast (left) and (B) fluorescence (right) images of PAO1 PvdA-eGFP grown in Succinate Media at 30°C for 48 h. These images correspond to a larger field of view of the images presented in Figure 2 (red selections). Scale bars = 2 µm.

(C) Jump-distance distribution (JD) representing the Euclidean distance travelled by ~5,500 PvdA-eYFP during a 16ms time interval. These data correspond to the JD observed in 11 cells measured in two independent experiments. (E) The corresponding empirical cumulative distribution function (ecdf) was fitted assuming a two-population diffusion model to retrieve diffusion coefficients of 0.06 [0.03 – 0.09] µm²/s and 0.48[0.46 – 0.51] µm²/s (median [IQR]) determined at 20°C for the restrained or bound (green) and mobile (orange) species, respectively – in very good agreement with observations made with PvdA-PamCherry.

Figure S3

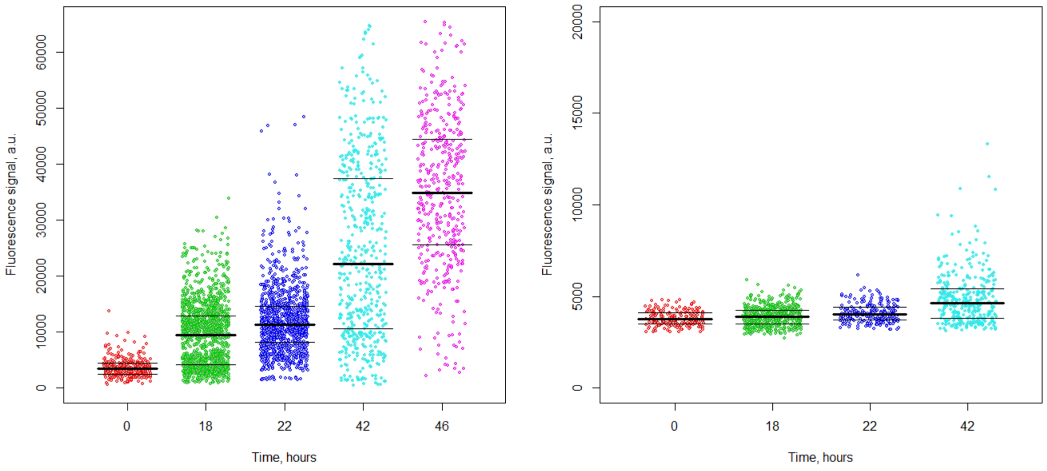


*Figure S3*

**Single-cell fluorescence signals of PvdA-eGFP (left) and PvdI-eGFP (right) measured at different cell growth time points after culture media was changed to succinate media.** Excitation wavelength was 488 nm. The fluorescence signal was filtered using a 488 nm long pass-filter. Each individual dot corresponds to the averaged fluorescence signal of one individual cell. Median and IQR intensity values of the cell signals are represented as horizontal lines. The level of expression of PvdA-eGFP was much higher than that of PvdI-eGFP in these conditions.

Figure S4

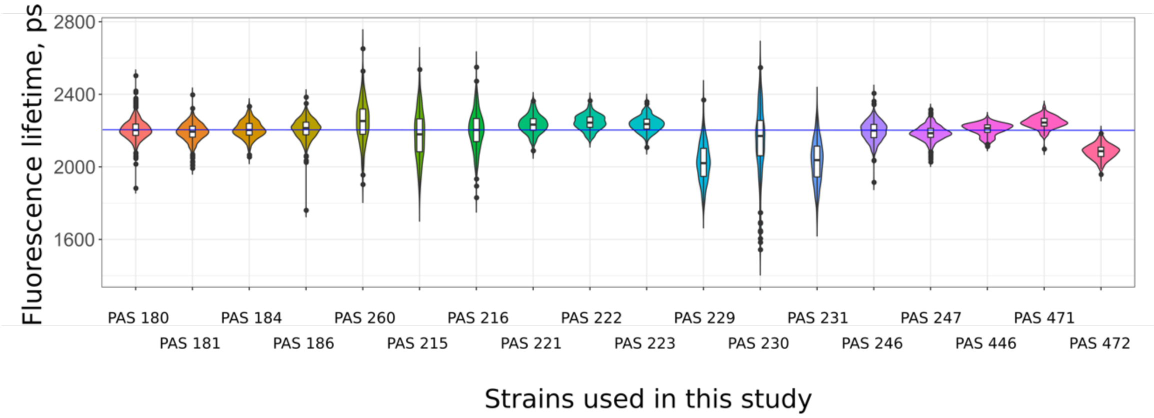


*Figure S4*

**Fluorescence lifetime distribution of the different strains used in this study.**
*Violin plot of the fluorescence lifetimes (one exponential model) for all the different strains used in this study.*

Figure S5

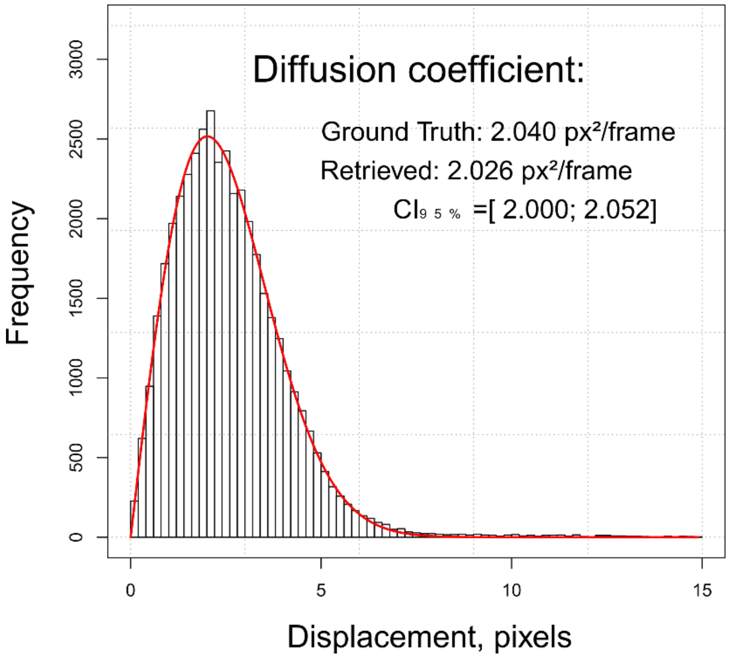


*Figure S5*

***Jump-distance distribution (JD) analysis of a simulated image data of Chenouard, N. et al.*** [5]

Data were extracted from the supplementary video 1 *corresponding to simulated vesicles diffusing according to a Brownian motion with a ground truth diffusion coefficient of 2.040px²/frame. The data were simulated at medium particle density and a signal-to-noise ratio of 4. To challenge the analysis pipeline fluorescent spots were tracked and analysed. The red line corresponds to the fit of the jump distances distribution. The estimation of the diffusion coefficient inferred from this data was 2.026 [ 2.000; 2.052 ] px²/frame, in excellent agreement with the ground truth.*

(4) Voisard, C.; Bull, C. T.; Keel, C.; Laville, J.; Maurhofer, M.; Schnider, U.; Dfago, G.; Haas, D. Biocontrol of Root Diseases ByPseudomonas Fluorescens CHA0: Current Concepts and Experimental Approaches. In *Molecular Ecology of Rhizosphere Microorganisms*; Wiley-VCH Verlag GmbH: Weinheim, Germany; pp 67–89.

(5) Chenouard, N.; Smal, I.; de Chaumont, F.; Maška, M.; Sbalzarini, I. F.; Gong, Y.; Cardinale, J.; Carthel, C.; Coraluppi, S.; Winter, M.; et al. Objective Comparison of Particle Tracking Methods. *Nat. Methods* 2014, *11* (3), 281–289.
